# Genome-wide mapping implicates R-loops in lineage-specific processes and formation of DNA break hotspots in neural stem/progenitor cells

**DOI:** 10.1101/2021.09.20.460546

**Authors:** Supawat Thongthip, Annika Carlson, Madzia P. Crossley, Bjoern Schwer

## Abstract

Recent work has revealed classes of recurrent DNA double-strand breaks (DSBs) in neural stem/progenitor cells, including transcription-associated, promoter-proximal breaks and recurrent DSB clusters in late-replicating, long neural genes. However, the mechanistic factors promoting these different classes of DSBs in neural stem/progenitor cells are not understood. Here, we elucidated the genome-wide landscape of DNA:RNA hybrid structures called “R-loops” in primary neural stem/progenitor cells in order to assess their contribution to the different classes of DNA break “hotspots”. We report that R-loops in neural stem/progenitor cells are associated primarily with transcribed regions that replicate early and genes that show GC skew in their promoter region. Surprisingly, the majority of genes with recurrent DSB clusters in long, neural genes does not show substantial R-loop accumulation. We implicate R-loops in promoter-proximal DNA break formation in highly transcribed, early replicating regions and conclude that R-loops are not a driver of recurrent double-strand break cluster formation in most long, neural genes. Together, our study provides an understanding of how R-loops may contribute to DNA break hotspots and affect lineage-specific processes in neural stem/progenitor cells.

## INTRODUCTION

Genome stability is important for cellular function but the genome of somatic cells shows much more plasticity than previously thought (Alt and Schwer, 2018). In mammals, somatic genomic alterations have traditionally been viewed primarily as a cause of cancer but they are emerging as drivers of organismal aging and brain disorders (Alt and Schwer, 2018; McKinnon, 2017; Schumacher et al., 2021; Wang and Lindahl, 2016; Wang et al., 2020). Somatic genomic alterations can arise from DNA double-strand breaks (DSBs) formed during normal cellular processes such as DNA replication and transcription. Mammalian cells use evolutionarily-conserved mechanisms to repair DSBs and maintain genome integrity (Alt and Schwer, 2018). In the nervous system, persistent DSBs caused by deficient repair can result in microcephaly, neurodegenerative disorders, and brain tumorigenesis (McKinnon, 2017).

Recent studies have identified several classes of recurrent DSBs in human and murine neural stem/progenitor cells (NSPCs) via high-throughput genome-wide translocation sequencing (HTGTS) (Schwer et al., 2016; Tena et al., 2020; Wang et al., 2020; Wei et al., 2018, 2016). Such classes include wide-spread, low-level DSBs, transcription start site (TSS)-proximal DSBs, and recurrent DSB clusters (RDCs) in long neural genes (Alt and Schwer, 2018; Schwer et al., 2016; Tena et al., 2020; Wang et al., 2020; Wei et al., 2018, 2016). Most RDCs in transcribed, long neural genes occur in gene bodies and are not associated with TSSs (Schwer et al., 2016; Tena et al., 2020; Wei et al., 2018, 2016), indicating that distinct mechanisms of DSB generation account for the different classes of recurrent DSBs in NSPCs.

Given the frequency and potential functional implications of RDCs in NSPCs, it is important to elucidate their mechanistic causes. In that regard, collisions of the transcription and replication machineries can cause genomic instability in mammalian cells (Hamperl and Cimprich, 2016). Indeed, prior studies of RDCs and copy number variations (CNVs) (Wang et al., 2020; Wei et al., 2016, 2016; Wilson et al., 2015) suggest that formation of the underlying DSBs may involve transcription/replication collisions in late-replicating regions (Alt and Schwer, 2018; Helmrich et al., 2011; Wilson et al., 2015). Consistent with that notion, RDCs in neural progenitor cells form within genes and are enhanced by DNA replication stress (Alt and Schwer, 2018; Wang et al., 2020; Wei et al., 2016). However, because the majority of long, transcribed, and late-replicating genes do not contain RDCs (Wei et al., 2016), additional factors must influence RDC formation.

To define such mechanistic factors, we considered whether transcription-related processes may affect RDC formation. Specifically, we asked whether RNA:DNA hybrid structures known as “R-loops” promote recurrent DNA breaks in NSPCs. R-loops consist of an RNA-DNA hybrid and the corresponding, displaced single-stranded DNA, thus forming a three-stranded nucleic acid structure (Crossley et al., 2019). Although R-loops have been known for over 50 years, their biological roles are still unclear. R-loops are emerging as important non-B DNA structures that form in transcribed loci (Crossley et al., 2019; Marnef and Legube, 2021). Traditionally, R-loops have been viewed as obstacles impeding ongoing transcription that need to be removed, and as sources of DNA damage that can induce single- and double-strand breaks and genomic instability (Costantino et al., 2014; Costantino and Koshland, 2015; Hamperl and Cimprich, 2014; Marnef and Legube, 2021; Skourti-Stathaki and Proudfoot, 2014).

How R-loops cause genomic instability is still unclear (Crossley et al., 2019). Pausing of transcription—which occurs when RNA polymerase II (RNAPII) progression is hindered—can induce RNAPII backtracking, which may form R-loops ahead of the backtracked RNAPII (Sheridan et al., 2019; Zatreanu et al., 2019). Thus, formation of R-loops could be an important contributing factor for the generation of both TSS-proximal DSBs and RDCs within long neural genes in NSPCs. To address this, we elucidated the genome-wide landscape of R-loop formation in NSPCs and assessed functional implications and relationships between these nucleic acid structures and classes of DSBs in NSPCs.

## RESULTS

### Genome-wide mapping of R-loops in NSPCs

Several classes of recurrent DSBs occur in mouse and human NSPCs, including DSB breakpoint clusters in long, transcribed and late-replicating genes and around transcriptional start sites (Schwer et al., 2016; Tena et al., 2020; Wang et al., 2020; Wei et al., 2018, 2016). To elucidate mechanistic factors involved in the formation of the different classes of recurrent DSBs, we assessed the genomic features of regions surrounding breakpoint junctions in NSPCs. We noted that the promoter regions—defined as regions ± 2 kb of the TSS—of active genes with HTGTS breakpoint junctions (Schwer et al., 2016; Wei et al., 2018, 2016) showed a significantly higher content of guanine/cytosine (GC) nucleotides (**Figure 1A**). This prompted us to consider the role of R-loops in the formation of recurrent DSBs in NSPCs, given that regions with high G density in the non-template strand are prone to R-loop formation (Hamperl and Cimprich, 2014). Moreover, R-loop formation has been implicated as a cause of genomic fragility in a subset of long human genes (Helmrich et al., 2011), suggesting a potential involvement in the formation of recurrent DSB clusters in long, neural genes (RDC-genes) (Tena et al., 2020; Wang et al., 2020; Wei et al., 2018, 2016).

**Figure 1.**
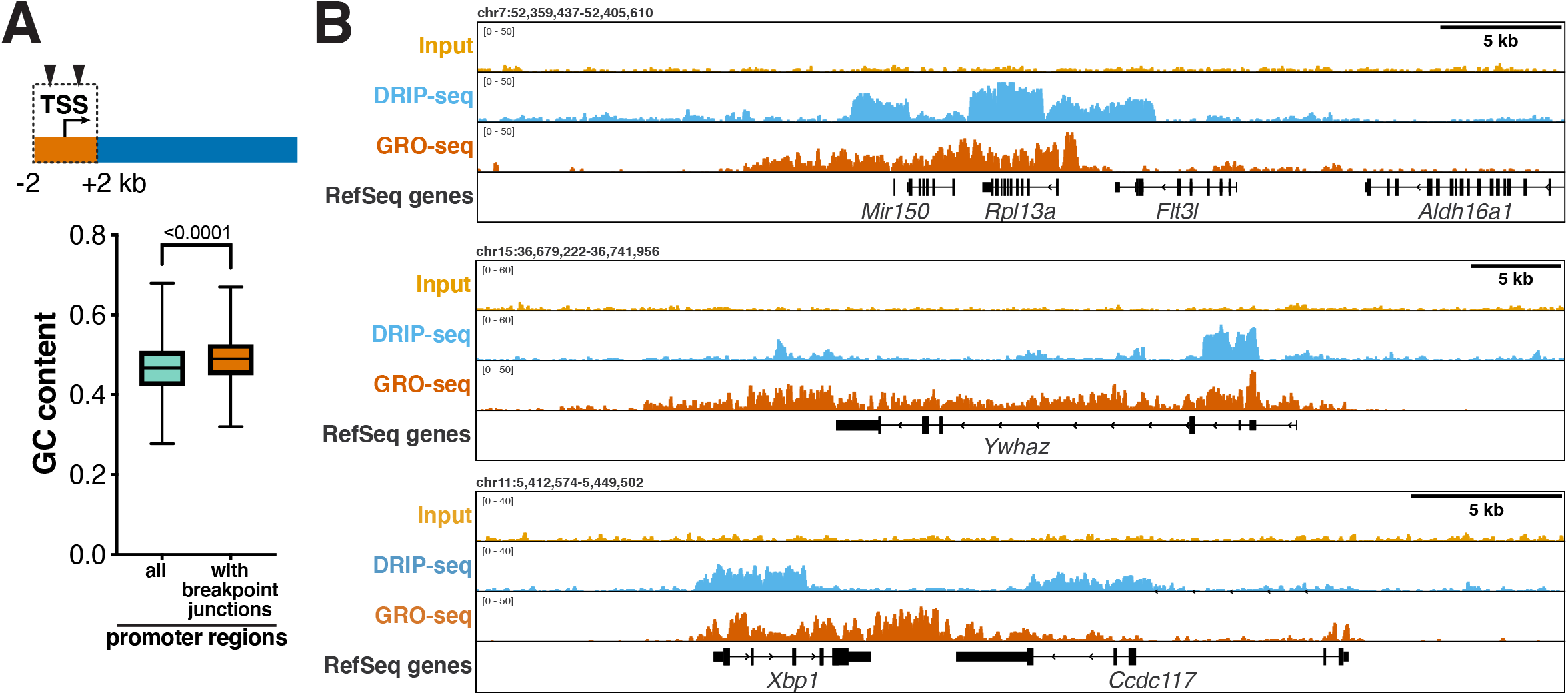
Elucidation of R-loops in neural/stem progenitor cells. **(A)** *Top*, illustration of genes with DNA breakpoints in the promoter region, defined as the 2-kb region surrounding the transcription start site (TSS). Triangles indicate HTGTS breakpoint junctions. *Bottom*, box-and-whiskers plot showing fractional GC content in promoter regions of all NCBI37/mm9 RefSeq genes and actively transcribed genes with HTGTS junctions within two kb of the TSS in NSPCs. Upper and lower box edges correspond to the 25^th^ and 75^th^ percentile; horizontal line indicates the median. *P* < 0.0001, Mann-Whitney U test. **(B)** Visualization of reads per kilobase per million (RPKM)-normalized DRIP-seq signal in input controls and DRIP samples over the indicated genomic regions. Combined signal from nine DRIP samples and matching input controls from three biological replicates is plotted. RPKM-normalized GRO-seq signal is plotted to show transcription. Location of RefSeq genes is shown.

To assess a potential role of R-loops in the formation of TSS-proximal DSBs or recurrent DSB clusters in long neural genes, we set out to elucidate the genomic landscape of R-loops in NSPCs. Recent high-throughput methods have enabled the genome-wide detection of R-loops (Chédin et al., 2021; Crossley et al., 2019; Marnef and Legube, 2021). Reliable, high-resolution mapping of R-loops has become possible by multiple approaches, including “DNA:RNA immunoprecipitation with deep sequencing” (DRIP-seq) (Chédin et al., 2021; Hamperl et al., 2017; Sanz et al., 2016; Stork et al., 2016). This approach relies on the S9.6 monoclonal antibody that specifically binds RNA:DNA hybrids and allows quantitative recovery of R-loops in conjunction with the high-resolution mapping capability of next-generation sequencing (Chédin et al., 2021).

We first validated the S9.6 antibody and DRIP approach by performing dot blots and DRIP-quantitative PCR (DRIP-qPCR) assays, using established positive and negative controls (**Figure S1A and B**). Next, we performed DRIP-seq in primary NSPCs isolated from postnatal day 7 mice in the presence of mild, aphidicolin-induced replication stress, as described (Schwer et al., 2016; Wei et al., 2016). To assess the quality of our R-loop mapping in NSPCs, we visualized raw DRIP-seq signal over the “gold standard” *Rpl13a* and the *Ywhaz* gene promoter region (Chédin et al., 2021). Consistent with previous reports in human cells (Chédin et al., 2021), we detected robust DRIP-seq signal over these regions in NSPCs (**Figure 1B**). Visual comparison of raw DRIP-seq signal in the mouse orthologs of human genes known to exhibit R-loops (Sanz et al., 2016; Stork et al., 2016) further confirmed the quality of our DRIP-seq analysis and revealed that R-loop formation in these genes is conserved between mice and humans and across cell types (**Figures 1B and S1C**). Analysis of NSPC DRIP-seq libraries from nine DRIP samples from three independent, biological replicates revealed a total of 22,132 R-loop peaks. R-loop peaks covered 1.01 ± 0.45 % (mean ± S.D.) of the NSPC genome, which is similar to the extent of R-loop formation reported for other cell types and species (Sanz et al., 2016).

### Comparative analysis of R-loop formation in NSPC and mouse embryonic stem cells

To compare our DRIP-seq results from NSPCs with published DRIP-seq data and gain insights into potential, lineage-specific features of R-loop formation, we obtained the deposited FASTQ files from DRIP-seq studies in pluripotent, mouse embryonic stem cells (ESCs) (Sanz et al., 2016). To enable direct comparisons, we performed all data analysis of NSPC and ESC DRIP-seq under identical conditions. R-loop peaks in NSPCs and ESCs showed a similar distribution across chromosomes (**Figure 2A**) and displayed a similar mean R-loop peak size of around 2 kb (NSPC, 2.19 ± 0.05 kb; ESC, 1.95 ± 0.02 kb; mean ± s.e.m) (**Figure 2B**). Overall, the ESC DRIP-seq data set contained a slightly higher total number of R-loop peaks (57,751) than detected in the combined NSPC DRIP-seq data, but when normalized via random down-sampling to the total peak number observed in NSPCs, ESCs and NSPCs showed similar absolute R-loop peak numbers and R-loop densities across chromosomes (**Figure S1D and E**).

**Figure 2.**
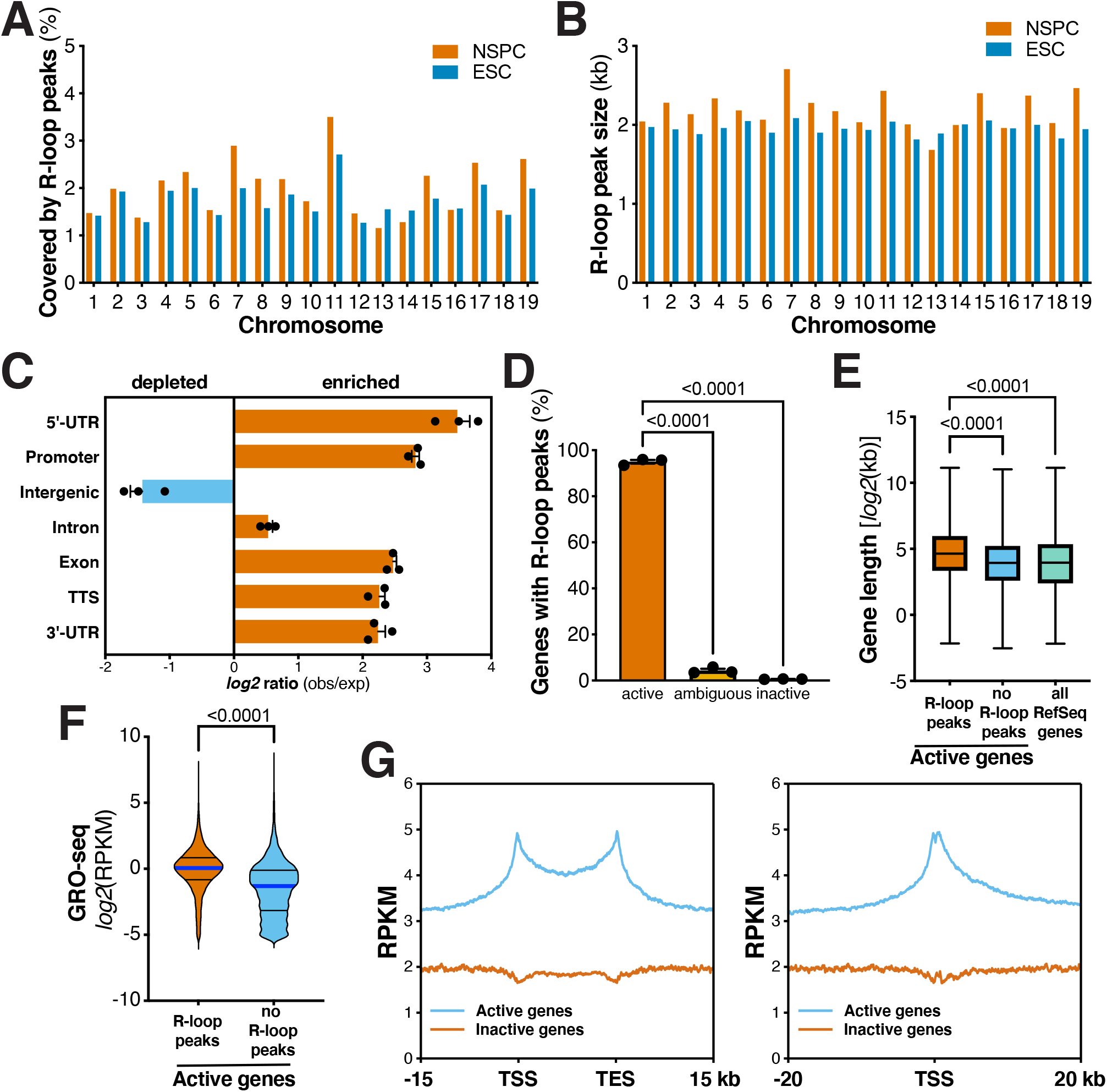
Genomic distribution of R-loops in neural stem/progenitor cells. **(A)** Coverage of the indicated chromosomes by R-loop peaks identified by custom Hidden Markov models (Sanz et al., 2016) in NSPCs and ESCs. Bar charts show mean coverage per chromosome from three biological replicates of DRIP-seq performed in NSPCs. **(B)** Mean R-loop peak size in NSPCs and ESCs, as in (A). **(C)** Genomic distribution of NSPC R-loop peaks across the indicated genome annotations compared to the expected genomic distribution. **(D)** Transcriptional status of genes with R-loop peaks in NSPCs. Data represent mean ± s.e.m. from three independent DRIP-seq experiments performed on biological replicates. *P*-values were determined by one-way ANOVA with Tukey’s *post hoc* correction for multiple comparisons. **(E)** Box-and-whiskers plot showing gene length of active genes with or without R-loop peaks in NSPCs and all RefSeq genes, for comparison. P-values were determined by one-way ANOVA with Tukey’s *post hoc* correction. **(F)** Transcriptional activity of active genes with or without R-loops in NSPCs. *P* < 0.0001, Mann-Whitney U test. **(G)** Metaplot analysis of RPKM-normalized raw DRIP-seq signal across active and inactive genes in NSPCs reveals enrichment at TSSs, gene bodies, and transcription end sites (TESs).

Further analysis revealed that R-loop peaks in NSPCs are significantly enriched in 5′-UTRs, promoters, introns, exons, transcription termination sites, and 3′-UTRs but are depleted in intergenic regions (**Figure 2C**). These patterns were consistent with the R-loop peak distribution in ESCs (**Figure S1F**). Overall, NSPC R-loop peaks were detected in 9,020 annotated genes (RefSeq NCBI37/mm9). Next, we assessed the transcriptional activity of genes containing R-loop peaks in NSPCs. Consistent with the notion that transcription promotes R-loop formation, the vast majority (99.17%) of genes containing R-loop peaks was either transcriptionally active (GRO-seq RPKM ≥0.025) or showed ambiguous (GRO-seq RPKM ≥0.0025 to <0.025) transcriptional activity. Only 75 (0.83%) of the 9,020 genes containing R-loop peaks in NSPCs were transcriptionally inactive (GRO-seq RPKM <0.0025) (**Figure 2D**). Analysis of ESC DRIP-seq data revealed that the vast majority (99.07%) of the 11,032 R-loop peak-containing ESC genes was either active or ambiguous—similar to the overall 99.17% of genes with R-loop peaks being either transcriptionally active or ambiguous in NSPCs (**Figure S1G**).

Surprisingly, active genes containing R-loop peaks in NSPCs were significantly longer than active genes without R-loops, with an average gene length of 62.68 ±1.39 kb (s.e.m.) and 38.7 ±0.98 kb (s.e.m.), respectively (**Figure 2E**). Consistent with this observation, R-loop peak-containing genes in ESCs were longer than genes without R-loops (54.58 ±0.98 kb vs. 36 ±1.26 kb; mean ±s.e.m.; **Figure S1H**). Moreover, active genes with R-loop peaks showed a significantly higher level of transcription than active genes without R-loop peaks in both NSPCs (**Figure 2F**) and ESCs (**Figure S1I**). These findings indicate that in both NSPCs and ESCs, R-loop-forming genes are generally longer and more actively transcribed than genes without R-loops.

To gain insights into the functions of R-loops and potential relationships to genomic stability in NSPCs, we further assessed the genomic distribution of DRIP-seq reads. DRIP-seq signal in NSPCs was present throughout the gene bodies of actively-transcribed genes and showed a robust enrichment around the transcription start sites (TSSs) and transcription end sites (TESs) of actively-transcribed genes (**Figure 2G**). These findings are consistent with the reported distribution of R-loops in other cell types and their involvement in regulatory functions in these regions (Ginno et al., 2013, 2012; Huertas and Aguilera, 2003; Promonet et al., 2020; Skourti-Stathaki et al., 2011; Stork et al., 2016).

### Factors promoting R-loop formation in NSPCs

To assess factors contributing to R-loop formation in NSPCs, we used a 4-state hidden-Markov model (StochHMM) (Ginno et al., 2013, 2012) to predict GC skew regions in the mouse genome. After identifying regions with GC skew, R-loop peak-containing genes in NSPCs were clustered into four skew classes (strong skew, weak skew, no skew, and reverse skew) (Ginno et al., 2013, 2012). This analysis revealed that the majority of R-loop-forming genes show GC skew, with only a minority (7.62%) exhibiting no GC skew (**Figure 3A**). Genes with R-loop peaks in ESCs showed a similar distribution across the four skew classes (**Figure 3A**), suggesting that GC skew is a universal feature of R-loop-forming genes in both cell types, consistent with findings in human cells (Ginno et al., 2013).

**Figure 3.**
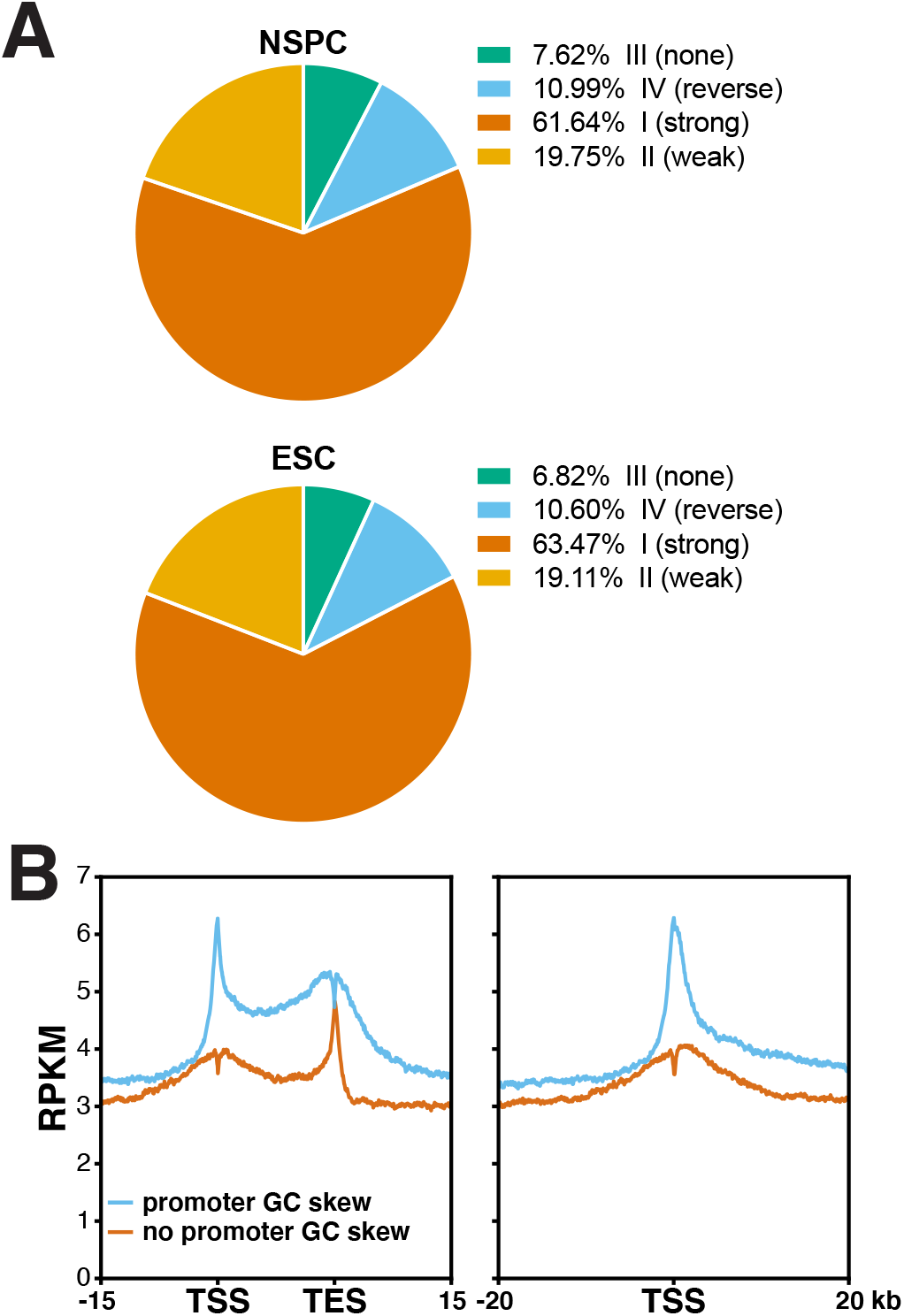
GC skew is a predictor of R-loop formation in NSPCs. **(A)** Pie charts showing distribution of genes with R-loop peaks in NSPCs (*top*) or ESCs (*bottom*) across the indicated classes of GC skew, showing that the majority of genes with R-loops exhibit GC skew. **(B)** Metaplot analysis of RPKM-normalized raw DRIP-seq signal across the TSS, gene body, and TES of active NSPC genes with or without GC skew in the promoter region. Fig.3A is high treshold. Fig.3B is low threshold

To further assess the impact of GC skew within the promoter region (±2kb of TSS) on R-loop formation in NSPC genes, we divided all 15,528 active genes into two groups; one group contained genes with GC skew within the promoter region (7,079 genes), whereas the other group contained genes without GC skew within the promoter region (8,449 genes). We then plotted the reads per kilobase per million (RPKM)-normalized DRIP-seq signal over these two groups of genes. Strikingly, genes with GC skew within 2 kb of the TSS showed a stronger DRIP-seq signal at the TSS and over the entire gene body and TES than genes without GC skew within the promoter region (**Figure 3B**). In contrast, the latter group of genes showed a robust peak at the TES (**Figure 3B**). These results suggest that GC skew in the promoter region is a strong predictor of overall R-loop formation throughout genes in NSPCs.

### Lineage-specific R-loop formation in ESCs and NSPCs

Next, we asked if R-loops in NSPCs are associated with genes involved in specific cellular functions and processes. To this end, we determined which genes show R-loop peaks in both ESCs and NSPCs (“common”), or uniquely in either cell type (“ESC unique”; “NSPC unique”) (**Figure 4**). The group of “common” R-loop genes contained 7,127 genes, representing 66.98% and 84.54% of active genes with R-loops in ESC and NSPCs, respectively. 1,303 genes (15.46%) were unique to NSPCs, and 3,514 genes (33.02%) were unique to ESCs (**Figure 4A**). **Figure 4B** shows examples, with *mitochondrial ribosomal protein 9* (*MrpS9*) being actively transcribed and forming R-loops in ESCs and *Pou3f3* (also known as *Brain-1*), a gene with roles in brain development (He et al., 1989) and intellectual disability (Snijders Blok et al., 2019) being unique to NSPCs. Core ESC transcriptional factors such as *Pou5f1, Lin28A* and *Zscan10*(*Zfp206*) were unique to ESCs (**Figure 4C**). *Pou3f2*(*Brain-2*), a gene involved in the establishment of neural cell lineage, neocortical development and associated with psychiatric disorders (C. Chen et al., 2018; Nakai et al., 1995) was unique to NSPCs (**Figure 4C**). Notably *Pou3f3/Brain-1* acts synergistically with *Sox11* and *Sox4* in neural development and we find that both show robust R-loop formation in NSPCs (**Figure S2A**). Moreover, R-loops in *Pou3f3/Brain-1* extended into the neighboring *Pantr1* (*Pou3f3 adjacent non-coding transcript 1*) gene, which encodes a long non-coding RNA implicated in glioma development (Guo et al., 2015).

**Figure 4.**
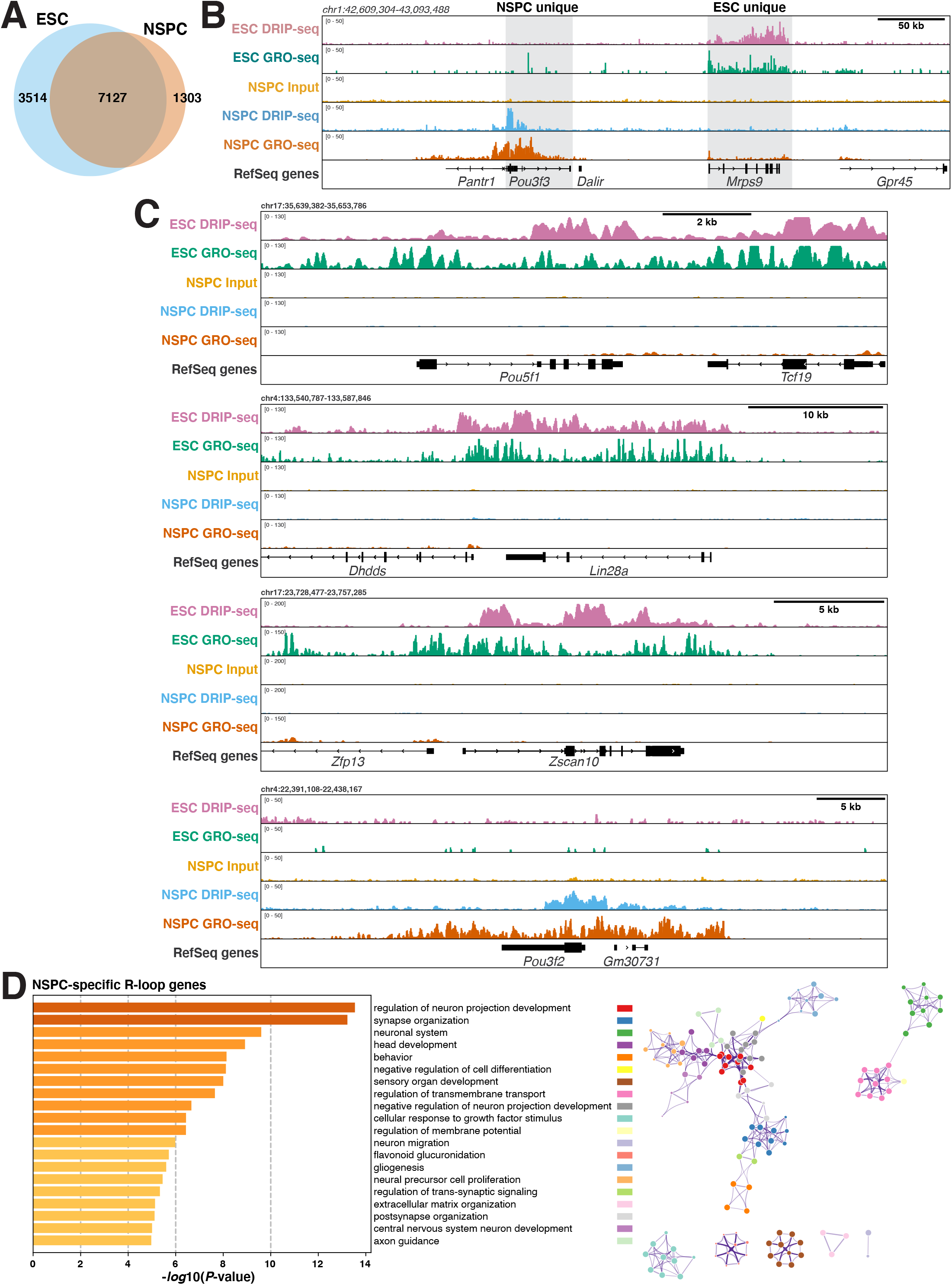
The landscape of R-loop formation in NSPCs suggests roles in lineage-specific processes. **(A)** Venn diagram showing the number of common and unique genes with R-loop peaks in ESCs and NSPCs. **(B)** Examples of genes forming R-loops in NSPCs or in ESCs. RPKM-normalized DRIP-seq and GRO-seq signals and RefSeq genes are shown. **(C)** Lineage-specific genes form R-loops in ESCs and NSPCs, illustrated as in (B). **(D)** *Left*, gene ontology (GO) analysis of NSPC-specific R-loop genes. Bars show significantly enriched GO terms and are colored by *P* values in *log* base 10. The Top 20 clusters are shown. *Right*, network visualization of the enriched terms shown on left, colored by cluster ID. Nodes sharing the same cluster ID are close to each other.

To assess the overall implications of R-loop formation within genes in the common, ESC-unique, and NSPC-unique sets, we performed pathway and process enrichment analyses (**Figures 4D and S2B-C**). Strikingly, we found that genes with unique R-loop formation in NSPCs were significantly enriched in processes related to neural development and function (**Figure 4D**). In stark contrast, shared R-loop genes were enriched for general biological processes (**Figure S2B**) and genes in the ESC-specific set showed enrichment of more general cellular processes, including DNA repair, cell cycle, and DNA replication (**Figure S2C**).

Together, these findings reveal potential lineage-specific function of R-loops and suggest that perturbation of R-loop formation in NSPCs may impact neural processes and development.

### Interplay between R-loop formation and DNA breakpoints in NSPCs

To determine an involvement of R-loop formation in the various classes of DSBs in NSPCs, we first examined the replication timing of R-loop peaks. R-loop peaks in NSPCs accumulated in early-replicating regions and showed, on average, a significantly earlier replication timing than Group 1-3 RDCs or the most robust 27 RDC-genes (Wei et al., 2018, 2016) (**Figure 5A**). These findings were corroborated in ESCs, where R-loop peaks showed a significantly earlier replication timing than the set of RDC candidates in ESCs (Tena et al., 2020) (**Figure S3A**). Notably, the replication timing of R-loop peaks in NSPCs and ESCs did not differ significantly (**Figure S3B**). These findings reveal that R-loop peaks preferentially occur in early-replicating regions of the genome in these two types of stem cells.

**Figure 5.**
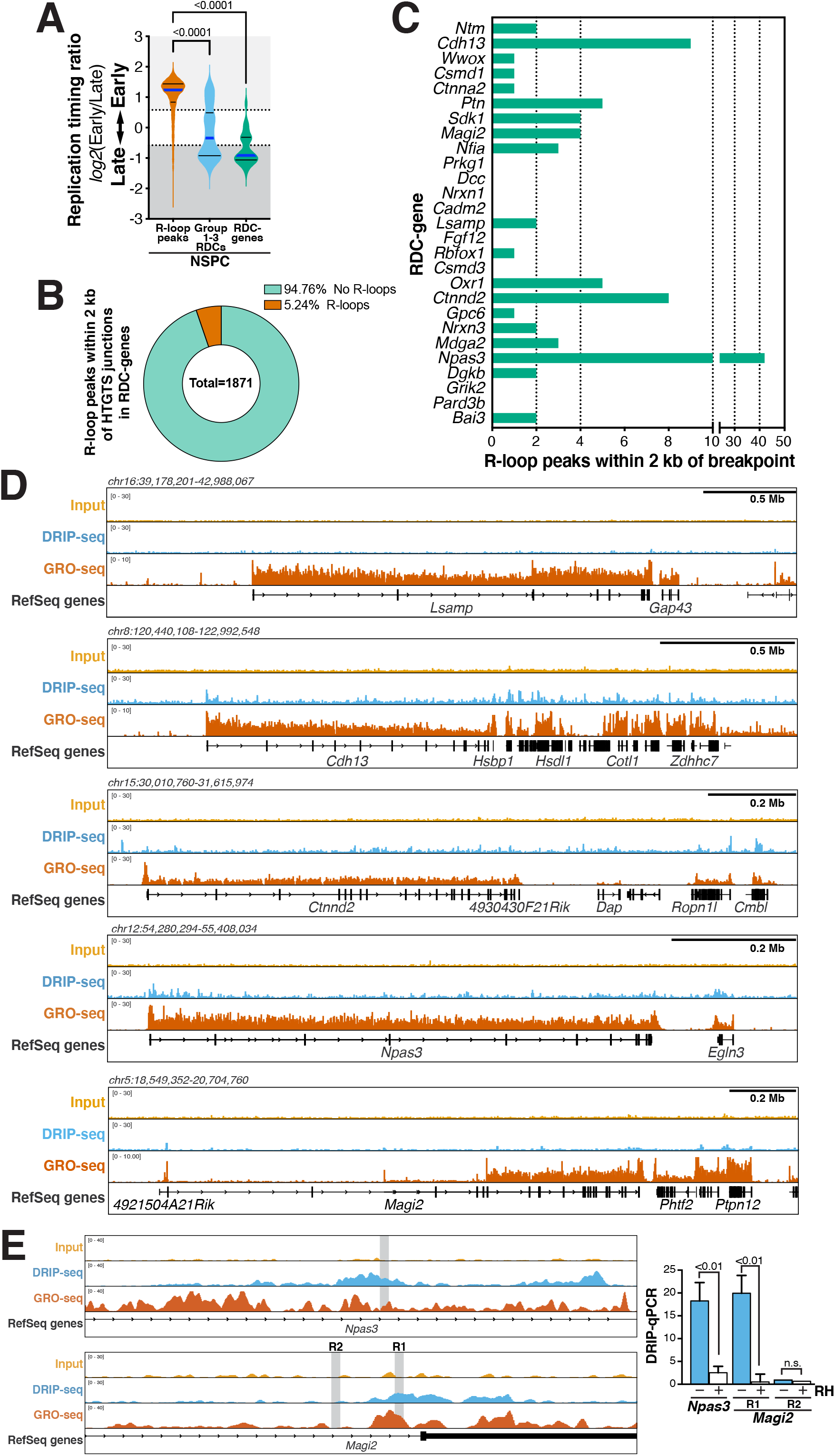
RDC-genes are not prone to extensive R-loop formation in NSPCs. **(A)** Violin plots showing the frequency distribution of replication timing ratios of all R-loop peaks, Group 1-3 RDCs, and the set of 27 RDC-genes in NSPCs. Median (blue line) and quartile lines (black) are shown. *P*-values were determined by one-way ANOVA with Tukey’s *post hoc* correction. **(B)** Parts-of-whole graph showing fraction of RDC-gene HTGTS junctions that fall within two kb of an R-loop peak. **(C)** Bar graphs indicating numbers of R-loop peaks within two kb of a breakpoint junction in RDC-genes in NSPCs. **(D)** RPKM-normalized DRIP-seq signal over the indicated RDC-genes. Combined signal from nine DRIP samples and matching input controls from three biological replicates of aphidicolin-treated NSPCs is plotted. RPKM-normalized NSPC GRO-seq signal is shown to indicate transcription. RefSeq genes are shown in black. **(E)** *Left*, zoomed-in visualization of R-loop signal in RDC-genes *Npas3* (*top*) and *Magi2* (*bottom*), as in (D). Grey rectangles indicate regions analyzed by DRIP-qPCR. *Right*, DRIP-qPCR analysis using primers over a region with (R1) or without (R2, negative control) an R-loop peak. Where indicated, samples were treated with RNase H prior to DRIP. Treatment with RNase H significantly suppressed the R-loop signal, consistent with R-loop formation in the tested regions. DRIP-qPCR signal intensity (mean ±s.e.m) shows fold enrichment over the *Snrpn* negative control region. *P*-values were determined by two-tailed, unpaired *t* test; n.s., not significant.

We further examined the formation of R-loops in the 27 RDC-genes (Wei et al., 2016). To this end, we determined the number of R-loop peaks within two kb of HTGTS breakpoint junctions (**Figure 5B**). Strikingly, within the most robust 27 RDC-genes, only 98 out of 1,871 breakpoint regions (5.24 %) contained an R-loop peak, whereas the vast majority (94.76%) of all RDC-gene breakpoint regions did not show R-loop formation (**Figure 5B**). To reveal potential differences in R-loop formation among the 27 RDC-genes, we determined the number of R-loop peaks within two kb of HTGTS breakpoints in each RDC-gene. Whereas some RDC-genes contained few to no R-loop peaks within two kb of an HTGTS junction, we noticed a range of R-loop formation, with *Npas3, Ctnnd2*, and *Cdh13* containing the most R-loop peaks (**Figure 5C-D**). DRIP-qPCR analysis confirmed the formation of R-loops (**Figure 5E**).

Based on these findings, we examined the overall correlation between HTGTS junction density and R-loop density across the RDC-genes. We found a moderate correlation (Spearman’s *r* = 0.4311, *P* = 0.0248) between R-loop peak density and HTGTS junction density (**Figure 6A**). Moreover, we found a moderate correlation (Spearman’s *r* = 0.5858, *P* = 0.0013) between GC content and R-loop peak density within RDC-genes (**Figure 6B**), suggesting that RDC-genes with higher GC content are more prone to R-loop formation.

**Figure 6.**
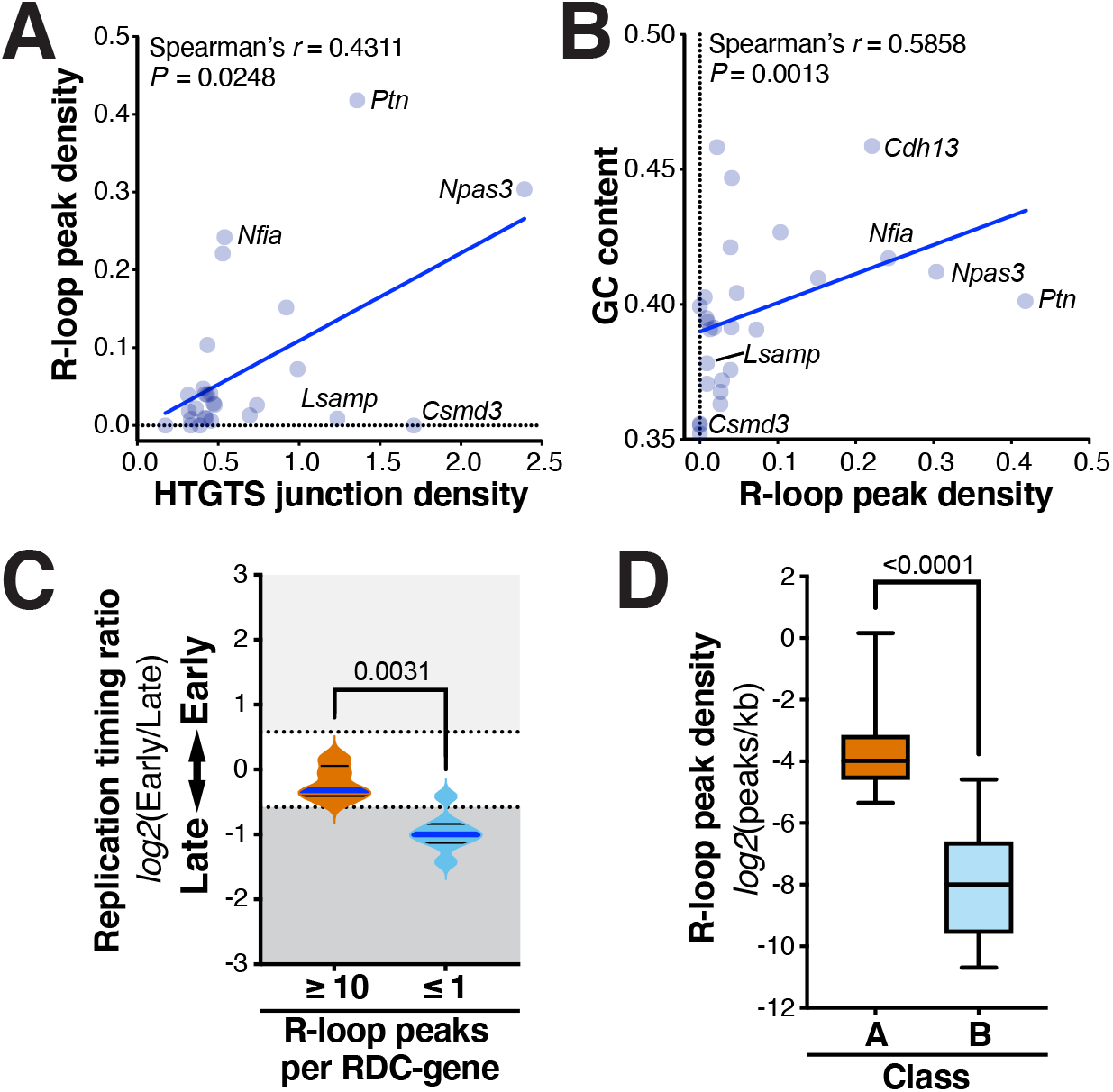
Characteristics of R-loop formation in NSPCs implicate R-loops in formation of TSS-proximal DSBs. **(A)** R-loop peak density and HTGTS junction density per 10 kb, respectively, are moderately correlated in RDC-genes, with some of the genes that show the most extensive RDCs (*e.g., Lsamp, Csmd3, Nrxn1*) showing low or absent R-loop peaks. **(B)** GC content and R-loop peak density across RDC-genes are correlated, indicating that the subset of RDC-genes with higher GC content forms more R-loops, which may contribute to at least some of the RDC formation in this set of genes. **(C)** RDC-genes with higher numbers of R-loop peaks show a significantly earlier replication timing than RDC-genes with low numbers of R-loop peaks. Violin plots show the frequency distribution of replication timing ratios in the two subsets of RDC-genes. Median (blue line) and quartile lines (black) are shown. **(D)** NSPC genes with TSS-proximal (±2 kb) breakpoint junctions (“A”) show a significantly higher R-loop peak density than RDC-genes (“B”) overall. *P*-values in (C) and (D) were determined by the Mann-Whitney U test.

RDCs in long neural genes tend to occur in large introns. Given the extensive splicing of transcripts of RDC-genes such as *Nrxn1* and *Nrxn3*, we hypothesized that co-transcriptional splicing of such large introns may make these genes prone to R-loop formation via reannealing of the nascent transcripts to the DNA. To our surprise, however, neither *Nrxn1* nor *Nrxn3* showed extensive R-loop formation, suggesting that RDCs in these long neural genes are not induced by a propensity for R-loop formation, even in the presence of mild replication stress. Moreover, in the latter context, we hypothesized that late-replication timing would promote R-loop formation via transcription/replication collisions. However, RDC-genes with the highest number of R-loop peaks showed significantly earlier replication timing than RDC-genes with the lowest number of R-loop peaks (**Figure 6C**). Together, these findings indicate that R-loops may contribute to DSBs in some RDC-genes. But surprisingly, we did not find abundant R-loop formation in the RDC-genes with the highest DSB density, suggesting that R-loops are not a major driver of RDC formation in these long, late-replicating genes.

We recently reported that breakpoint junctions are enriched around active TSSs in NSPCs (Schwer et al., 2016). To assess a potential role of R-loops in promoting this class of DSBs, we compared the gene length-normalized R-loop peak density of active genes of average length (*i.e*., 5.49–25.49 kb) containing an HTGTS junctions within two kb of the TSS (“Class A”) and those of the most robust RDC-genes containing at least one R-loop peak (“Class B”), respectively (as shown in **Figure 5C**, we did not detect any R-loop peaks in *Csmd3, Cadm2, Dcc, Grik2, Fgf12, Nrxn1, Pard3b*, and *Prkg1*). Actively-transcribed NSPC genes of average gene length with TSS-proximal breakpoint junctions displayed a significantly higher R-loop peak density than RDC-genes (**Figure 6D**). These findings further suggest that R-loops are not a major contributor to breakpoints in RDC-genes and implicate R-loops in the formation of promoter-proximal DSBs. Together, our findings support the notion that the classes of recurrent DSBs in NSPCs, *i.e*., TSS-associated DSBs and DSBs in the gene bodies of long, transcribed neural genes, are caused by different mechanisms, with the former class being promoted by R-loop formation.

## DISCUSSION

In recent work, several classes of recurrent DSBs have been discovered in neural progenitor cells (Schwer et al., 2016; Wang et al., 2020; Wei et al., 2018, 2016), suggesting that somatic brain mosaicism may arise at the level of rapidly dividing NSPCs during fetal brain development. However, the underlying mechanistic causes are unclear. We initially hypothesized that R-loops contribute to RDC formation in long neural genes based on several considerations: (*1*) R-loops can form at sites of RNA polymerase pausing caused by transcription-replication machinery collisions (Crossley et al., 2019; Hamperl et al., 2017; Huertas and Aguilera, 2003); (*2*) long neural genes that form RDCs undergo extensive co-transcriptional splicing and pre-mRNA processing (Ameur et al., 2011), which can induce DSBs via R-loop formation (Li and Manley, 2005); and (*3*) earlier work has implicated R-loop formation in common fragile site formation within a subset of large human genes, based on slot blot hybridization experiments in the *FHIT* locus (Helmrich et al., 2011). Thus, we were surprised to find that R-loops do not preferentially and robustly form in RDCs-genes that undergo extensive splicing—such as *Nrxn1* and *Nrnx3*— nor in *Lsamp* (**Figure 5C-D**), one of the most robust RDCs in mouse NSPCs and human cells (Wei et al., 2016; Wilson et al., 2015).

There are several—not mutually exclusive—potential explanations for our findings: R-loop formation in RDC-genes such as *Lsamp, Cadm2, Nrxn1, Csmd3*, and others may be highly dynamic and transient, thus making these R-loops difficult to capture in primary NSPCs. However, this seems unlikely as we detected robust R-loop signal in promoter regions and TESs where R-loops are known to assemble in a dynamic and transient manner. A more likely interpretation is that RDC formation in long genes such as *Lsamp, Csmd3, Nrxn1, Nrxn3, Csmd3*, and others is not primarily driven by R-loop formation. This conclusion is further supported by the generally lower GC content of RDC-genes (**Figure S3C**). Moreover, our findings are consistent with recent work showing a paucity of R-loops in the center of large human genes and proposing that the determining factor of replication stress-induced genomic instability is transcription-dependent persistence of unreplicated DNA into mitosis rather than R-loop formation (Park et al., 2021).

We note, however, that there are differences in the extent of R-loop formation among the group of long RDC-genes in NSPCs. For example, the RDC-genes *Cdh13, Npas3* and others (**Figure 5C-D**) showed varying degrees of R-loop formation, which may contribute to DNA breakpoint formation in this subset of RDC-genes. Notably, this latter set of RDC-genes tends to show earlier replication timing and higher GC content than RDC-genes without R-loops (**Figure 6B-C**). The observed differences in R-loop formation among RDC-genes in NSPCs may be due to differential enrichment of factors that affect the formation or removal of R-loops, which warrants further investigation. Together, our findings on R-loop formation in NSPCs undergoing mild replication stress suggest that factors such as unique topological stresses or other factors (Helmrich et al., 2013), rather than extensive R-loop accumulation *per se*, promote DSB formation in the class of late-replicating RDCs that overlaps with the gene bodies of large genes. Further studies of these genes are clearly important and may reveal the precise mechanisms by which these regions undergo extensive DSB formation.

Prior work indicated that the mechanisms promoting TSS-proximal DSBs and DSBs in the gene bodies of long genes are distinct (Alt and Schwer, 2018; Schwer et al., 2016; Wei et al., 2016). Our results implicate R-loops in the formation of DNA breaks in the promoter-proximal regions of genes in NSPCs. Specifically, NSPC genes with breakpoint junctions within two kb of the TSS show significantly higher R-loop peak density than RDC-genes (**Figure 6D**), consistent with the notion that transcription-associated processes contribute to promoter-proximal DSBs in NSPCs (Schwer et al., 2016). Further work is required to define the molecular factors responsible for DSB formation in R-loop-prone regions in the NSPC genome.

Importantly, R-loops may—at least in some contexts—function as “beneficial” structures that help with DSB repair (Marnef and Legube, 2021). This latter notion is based on observations that R-loops form at DSB sites in response to various types of DNA damage, and DSB-induced R-loops may form in *cis* as a response to DSB-mediated repression of transcription (Marnef and Legube, 2021). Future work will need to address whether some of the R-loops we observe in the vicinity of DNA breakpoints could have such roles in DSB repair within R-loop forming regions in NSPCs, for example by recruiting repair factors such as Rad52.

Our findings demonstrate that primary NSPCs show R-loop enrichment at TSSs and TESs (**Figure 2G**), consistent with a role of R-loops in regulation of gene expression and transcription termination (Arab et al., 2019; Crossley et al., 2019; Ginno et al., 2013; Skourti-Stathaki et al., 2011). We found that NSPCs share a common set of R-loop containing genes with ESCs but also contain a substantial fraction of unique R-loop genes (**Figure 4A**). These latter findings indicate a cell lineage-specific role of R-loops. Indeed, we found that NSPC-specific R-loop genes are significantly enriched in genes with functions in neural development and neural function (**Figure 4D**).

Further studies of R-loop biology in neural progenitors may reveal important insights into processes ranging from neurodevelopment to neurological diseases. Factors modulating R-loop formation may play roles in the generation of somatic alterations during neurodevelopment, which may affect the extent of somatic brain mosaicism and occurrence of brain disorders. In the latter context—for reasons that are unclear—mutations in R-loop processing factors cause neurological disease in humans. Based on our finding of R-loop formation in genes with neural functions, we speculate that R-loop-mediated neurological disorders may have a previously unanticipated neurodevelopmental etiology that originates at the level of neural progenitors—for example by affecting genomic stability or function of epigenetic R-loop readers (Arab et al., 2019). On a related note, DNA damage caused by augmented R-loop formation has been proposed as a unifying mechanism for myelodysplastic syndromes induced by splicing factor mutations (L. Chen et al., 2018). Based on our work here, we hypothesize that splicing factor mutations may promote neurological disorders via increased R-loop formation and DNA damage in neural progenitors.

Of note, R-loops have been suggested to promote the instability of pathogenic repeat sequences in trinucleotide repeat-associated neurological diseases such as Huntington’s disease (Lin et al., 2010; McIvor et al., 2010; Reddy et al., 2011; Richard and Manley, 2017). Huntingtin (*Htt*) can form R-loops when transcribed *in vitro* (Lin et al., 2010; McIvor et al., 2010; Reddy et al., 2011; Richard and Manley, 2017). Our DRIP-seq analysis reveals that *Htt* forms R-loops *in vivo* in NSPCs (**Figure S4A**), suggesting potential contributions of R-loops to Huntington’s disease pathology.

Together, our comparative analysis of DSB breakpoint junctions and R-loop formation in NSPCs supports the notion that R-loops can cause DNA damage and mutagenesis, two hallmarks of cancer (Crossley et al., 2019; Groh and Gromak, 2014; Hanahan and Weinberg, 2011). Notably, R-loops have recently been implicated in the etiology of *Embryonal Tumor with Multilayered Rosettes*, a malignant brain tumor almost exclusively affecting young children, via induction of DNA breaks (Lambo et al., 2019). Our findings that R-loops associate with DSBs in NSPCs may thus suggest a role in the etiology of brain tumors more broadly. Indeed, several of the R-loop-forming genes we identified in NSPCs show rearrangements and mutations in human low-grade and high-grade gliomas, including *Raf1, Daxx, Fgfr1, Lztr1, and H3F3A* **(Figure S4B)** (Frattini et al., 2013; Johnson et al., 2017; Liu et al., 2014), which warrants further studies of the role of R-loops in brain cancer development.

## EXPERIMENTAL PROCEDURES

### Anti-DNA:RNA hybrid S9.6 antibody

Hybridoma cells producing the monoclonal S9.6 antibody were obtained from ATCC (HB-8730; RRID:CVCL_G144) and grown in chemically-defined, protein-free CD hybridoma medium (Gibco 11279023). S9.6 antibodies were purified according to standard procedures (Stork et al., 2016) by using a HiTrap Protein G HP column (GE Healthcare), followed by extensive washing with 20 column volumes of PBS and elution with five column volumes of elution buffer (0.1 M glycine-HCl, pH 2.7). Antibody-containing fractions were assessed for purity by SDS-PAGE and Coomassie blue staining and sequentially dialyzed against PBS and 50% (v/v) glycerol/PBS. Antibody concentration was adjusted to 1 mg/mL. Purified S9.6 antibodies were validated by dot blot and DRIP-qPCR analysis along with commercially available S9.6 antibodies (Kerafast, ENH001; S.H. Leppla, NIH).

### Culture of primary NSPCs

Frontal brains of postnatal day 7 mice were used to isolate and culture NSPCs as described (Wei et al., 2016). All experiments were authorized by the Institutional Animal Care and Use Committee and Institutional Biosafety Committee at the University of California, San Francisco (Protocol AN182936). Where indicated, cells were treated with 0.5 μM aphidicolin (Sigma, A4487) for 48 h before processing for DRIP.

### DRIP analysis and genome-wide R-loop mapping by DRIP-seq

Genomic DNA for DRIP was isolated and digested with *EcoRI, HindIII, BsrgI, SspI, XbaI* (all from NEB) at 37 °C, as described (Ginno et al., 2012; Stork et al., 2016). RNase A treatment prior to DRIP was performed as described (Hamperl et al., 2017; Sanz et al., 2016). For Ribonuclease H (RNase H) treatment, 8 μg of DNA were treated with 30 U of RNase H (NEB, M0297) for 16 h at 37 °C. Digested DNA was phenol/chloroform-extracted, precipitated, washed and resuspended as described (Stork et al., 2016). For each DRIP reaction, 4.4 μg DNA were incubated with 10 μg S9.6 antibody for 16 h at 4 °C (Stork et al., 2016), followed by incubation with magnetic protein G beads (Dynabeads, ThermoFisher Scientific) for 2 h at 4 °C. Samples were washed for 3 × 10 min with 140 mM NaCl, 0.05% (w/v) Triton X-100, 10 mM NaPO4, pH 7.0, at room temperature and eluted in 10 mM EDTA, 0.5% (w/v) SDS, 50 mM Tris-Cl, pH 8.0 containing Proteinase K for 45 min at 55 °C (Stork et al., 2016). DNA was phenol/chloroform-extracted and precipitated as described above. Per biological replicate, three repeat DRIP reactions were first analyzed separately by DRIP-qPCR to verify the DRIP procedure, as described (Sanz and Chédin, 2019). Three DRIP reactions per biological replicate were then pooled for preparation of each DRIP-seq library. Input and DRIP DNA was sonicated (Diagenode Bioruptor) to a size of *300–700 bp and DRIP-seq library preparation was performed as described (Ginno et al., 2012; Stork et al., 2016), using NEB E6050 for end repair; NEB M0212 for A-tailing, and NEB E7335 for adapter ligation. 12 cycles of PCR were performed for library amplification and libraries were cleaned and size selected (Ampure XP beads; A63880, Beckman Coulter) as described (Sanz and Chédin, 2019). Libraries were assessed and quantified by using the Qubit HS assay (Invitrogen Thermo Scientific), Bioanalyzer High Sensitivity DNA Analysis (Agilent), and qPCR-based KAPA Library Quantification (KK4824, Kapa Biosystems). Pooled libraries were sequenced on the Illumina HiSeq next-generation sequencing platform.

### S9.6 dot blot analysis

DNA (5’-GTTCCCATATCCCGGACGAGCCC-3’) and RNA oligonucleotides (5’-rGrGrGrCrUrCrGrUrCrCrGrGrGrArUrArUrGrGrGrArArC-3’) were annealed in RNA:DNA hybridization buffer (20 mM NaCl, 10 mM Tris-HCl, pH 8.0) and spotted onto Nylon membranes (GE Healtcare, RPN303B). Membranes were dried, UV crosslinked (120 mJ/cm^2^), incubated in blocking buffer (10% (w/v) non-fat dry milk in TBS with 0.1% (w/v) Tween-20) and probed with 0.5 μg/mL S9.6 antibody in blocking buffer for 1 h at room temperature. After three washes in TBS with 0.1% (w/v) Tween-20, membranes were incubated with secondary anti-mouse-HRP antibodies in blocking buffer for 1 h at room temperature, washed again, developed (ECL), and exposed to film.

### Bioinformatic and statistical analysis

DRIP-seq and GRO-seq reads were adapter trimmed (TrimGalore 0.6.6; https://github.com/FelixKrueger/TrimGalore), aligned to the NCBI37/mm9 genome, and processed as described (Sanz et al., 2016; Wei et al., 2016). Mouse ESC GRO-seq and DRIP-seq FASTQ files were obtained via GEO (Min et al., 2011; Sanz et al., 2016) and processed in parallel with NSPC GRO-seq (Wei et al., 2016) and NSPC DRIP-seq data. Duplicate reads were removed during processing. We used a Hidden Markov Model-based peak calling algorithm for identification of DRIP-seq peaks, as described (Sanz et al., 2016). Normalized genome-wide densities of uniquely mapped reads were determined by deepTools2.0 (Ramírez et al., 2016). Read densities were assessed via bedtools coverage and normalized to total mapped reads per sample. Reads per kb per million (RPKM)-normalized bigWig tracks were generated from BAM files containing uniquely mapped reads using deepTools2.0 and visualized in IGV (Robinson et al., 2011). Nucleotide content was analyzed by bedtools version 2.29.2 (Quinlan and Hall, 2010). Genome annotations were determined by HOMER version 4.11.1 (Heinz et al., 2010). HTGTS breakpoint junction data was analyzed as described (Schwer et al., 2016; Tena et al., 2020; Wei et al., 2018, 2016). Median replication timing ratios were determined using Repli-chip data (Hiratani et al., 2008; Weddington et al., 2008) and custom Python scripts, as described (Wei et al., 2016). Statistical analysis was performed in GraphPad Prism 9.2.0 and in R (R Core Team, 2021).

### Identification of genomic regions with GC skew

We applied the two-phase SkewR pipeline 1.00b (Ginno et al., 2013, 2012) developed to define regions displaying GC skew in the human genome to the mouse NCBI37/mm9 genome. SkewR uses a four-state hidden-Markov model (StochHMM) to predict GC skew regions (Ginno et al., 2013, 2012). The algorithm involves training on regions from verified R-loop-forming regions in human and mouse regions. Regions displaying GC skew were identified and genes were clustered into four skew classes (strong skew, weak skew, no skew, and reverse skew) by using the most stringent threshold model file (GC_SKEW_1mil.hmm) (Ginno et al., 2013, 2012). For metagene plots, the GC_SKEW_7600 model file was used (Ginno et al., 2013).

### Pathway and process enrichment analysis of R-loop-containing genes

For each gene list, pathway and process enrichment was analyzed using the GO Biological Processes, KEGG Pathway, Reactome Gene Sets, CORUM, TRRUST, PaGenBase, WikiPathways and PANTHER Pathway ontology sources and Metascape version 3.5.202108015 (Zhou et al., 2019). All mouse genes were used as the enrichment background. Enrichment terms with P < 0.01, at least three counts, and an enrichment factor (counts observed vs. counts expected by chance) of > 1.5 were identified and clustered based on their similarities. P-values were calculated based on the accumulative hypergeometric distribution. q-values were calculated by the Benjamini-Hochberg procedure to account for multiple testing. Kappa scores were used as the similarity metric for hierarchical clustering of enriched terms and sub-trees showing > 0.3 similarity were considered a cluster, with the most statistically significant term chosen to represent the cluster (Zhou et al., 2019). Clusters were visualized by Cytoscape (Shannon et al., 2003).

## AUTHOR CONTRIBUTIONS

B.S. designed and planned the study; S.T., A.C., and B.S. designed experiments and performed research; M.P.C. developed experimental approaches; S.T., A.C., and B.S. analyzed data; B.S. supervised the research and wrote the manuscript. All authors commented on the manuscript.

## ACKNOWLEDGMENTS

We thank Dr. K. Cimprich for helpful discussions and Drs. P. Oberdoerffer and Jeongkyu Kim for sharing protocols. We thank members of the Schwer lab, Dr. Z. Yang, and Dr. T. E. Wilson for critical reading of the manuscript. This work was supported by the UCSF Brain Tumor SPORE Career Development Program, the American Cancer Society, the UCSF Program for Breakthrough Biomedical Research (which is partially funded by the Sandler Foundation), the Shurl and Kay Curci Foundation, and NIH R01 AG064363 (to B.S.). B.S. is a Kimmel Scholar of The Sidney Kimmel Foundation, is supported by a Carol and Gene Ludwig Award for Early Career Research, and holds the Suzanne Marie Haderle and Robert Vincent Haderle Endowed Chair at UCSF.

## SUPPLEMENTAL FIGURE LEGENDS

**Figure S1.**
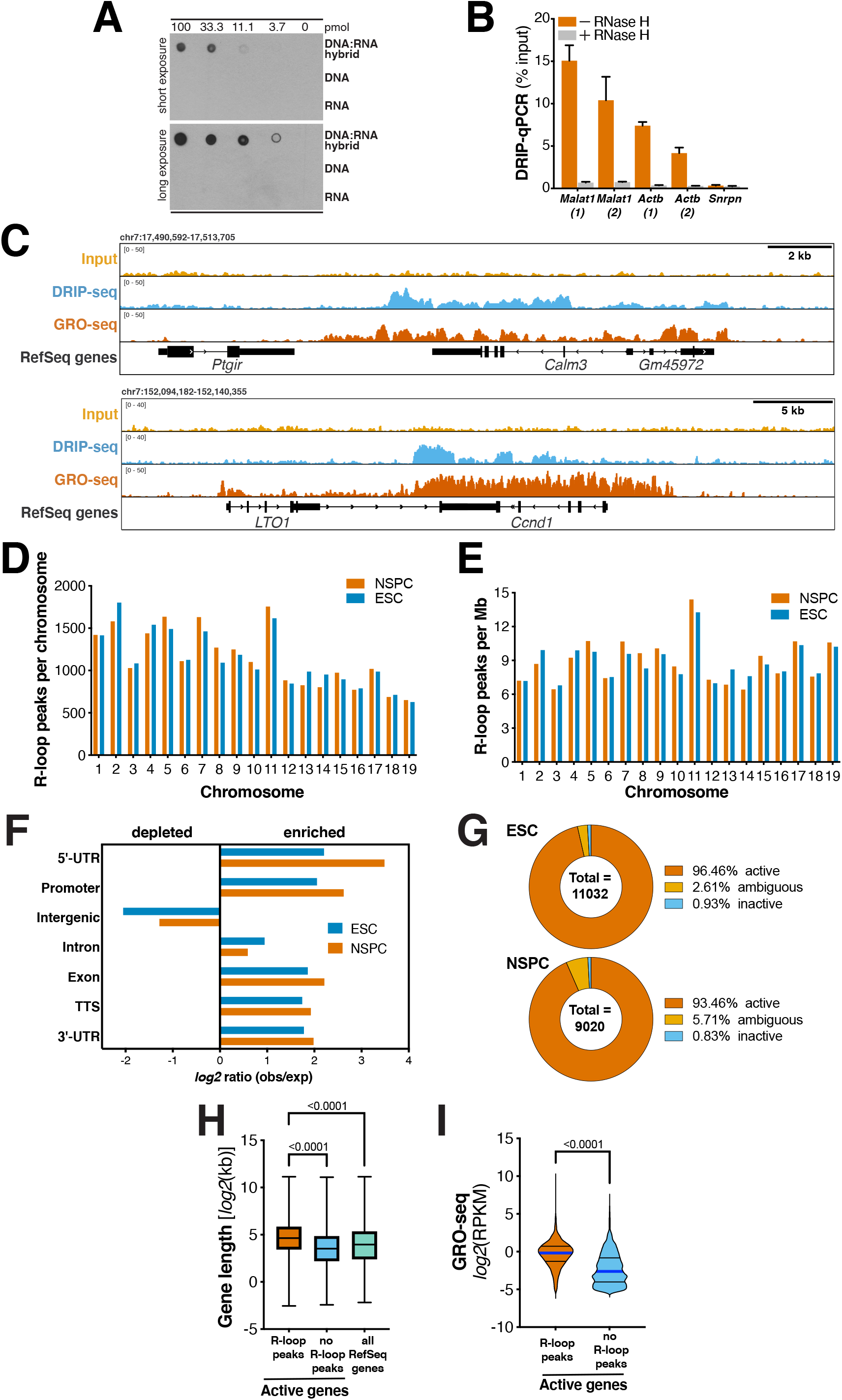
DRIP-seq analysis of NSPCs and comparison with ESCs. **(A)** Dot blot analysis of serial dilutions of *in vitro* synthesized DNA:RNA hybrids (*top*) or equivalent amounts of the corresponding DNA (*middle*) or RNA (*bottom*) with the S9.6 anti-DNA:RNA hybrid antibody. **(B)** DRIP-qPCR analysis of the indicated positive and negative control regions in NSPCs. Bars show DRIP-qPCR signal as percentage of input. Pre-treatment with RNase H abolished the DRIP-qPCR signal. Data represent mean ±s.e.m. from three DRIP reactions performed in NSPCs. **(C)** RPKM-normalized NSPC DRIP-seq signal over the indicated genes, which also form R-loops in human cells (Sanz et al., 2016; Stork et al., 2016). Combined signal from nine DRIP samples and matching input controls from three biological replicates of APH-treated NSPCs is plotted. RPKM-normalized NSPC GRO-seq signal shows transcriptional activity. RefSeq genes are shown in black. **(D)** Absolute numbers of R-loop peaks per indicated chromosome in NSPCs (mean from three biological replicates) and ESCs. **(E)** R-loop peak density, shown as in (D). **(F)** Comparison of distribution of R-loop peaks across the shown genome annotations in NSPCs and ESCs. **(G)** Parts-of-whole graphs showing transcriptional activity of R-loop peak-containing genes in ESCs (*top*) and NSPCs (*bottom*), as determined by GRO-seq analysis. **(H)** Box-and-whiskers plot showing gene length of active genes with or without R-loop peaks in ESCs and all RefSeq genes, for comparison. *P*-values were determined by one-way ANOVA with Tukey’s *post hoc* correction. **(I)** Transcriptional activity of active genes with or without R-loops in ESCs. *P* < 0.0001, Mann-Whitney U test.

**Figure S2.**
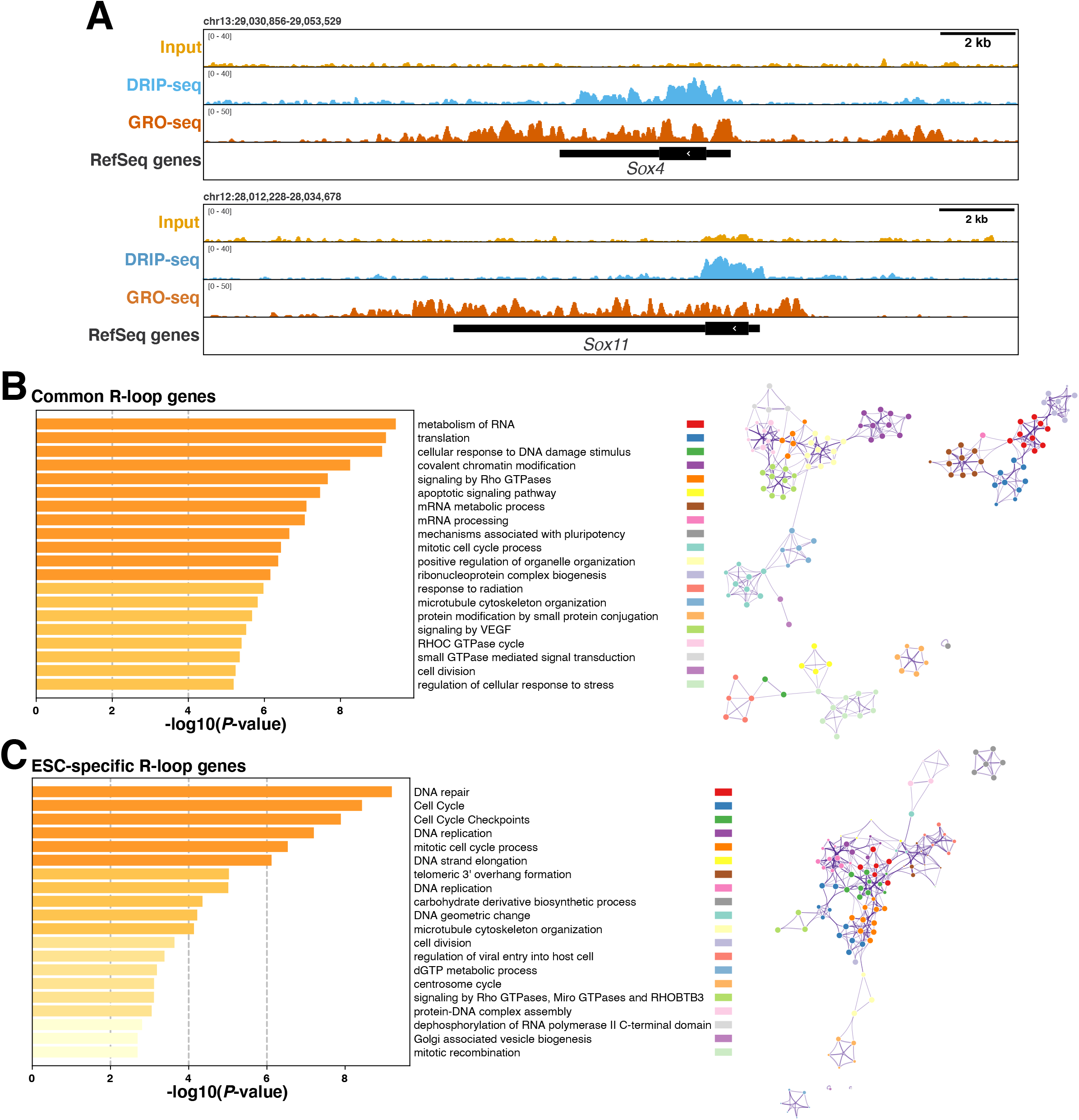
R-loop formation in genes with general and lineage-specific functions in NSPCs and ESCs. **(A)** R-loop formation in *Sox4* and *Sox11* in NSPCs. RPKM-normalized NSPC DRIP-seq signal is shown. **(B)** *Left*, GO analysis of genes showing R-loop peaks in both NSPCs and ESCs. Bars show significantly enriched GO terms and are colored by *P* values in *log* base 10. The Top 20 clusters are shown. *Right*, network visualization of the enriched terms shown on left, colored by cluster ID. Nodes sharing the same cluster ID are close to each other. **(C)** *Left*, GO analysis of genes showing R-loop peaks only in ESCs, as in (B).

**Figure S3.**
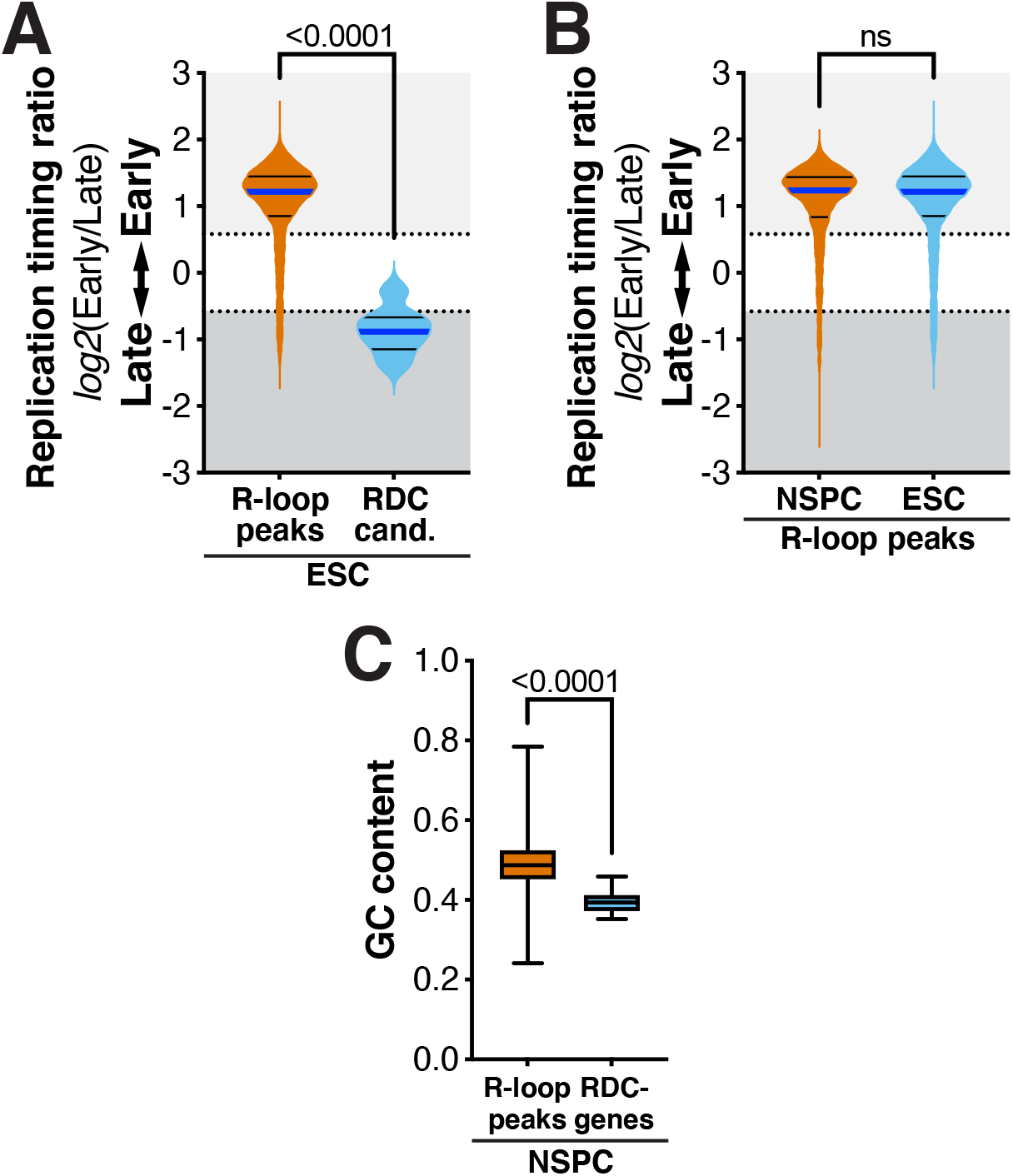
Replication timing profiles and GC content of R-loop peaks and RDCs. **(A)** Violin plots showing the frequency distribution of replication timing ratios of all R-loop peaks and the set of RDC candidates in ESCs (Tena et al., 2020). **(B)** Comparison of replication timing of all R-loop peaks in NSPCs and ESCs. Median (blue line) and quartile lines (black) are shown. **(C)** Box-and-whiskers plot showing fractional GC content of all R-loop peaks and RDC-genes in NSPCs. Upper and lower box edges correspond to the 25^th^ and 75^th^ percentile; horizontal line indicates the median. *P*-values in (A–C) were determined by the Mann-Whitney U test.

**Figure S4.**
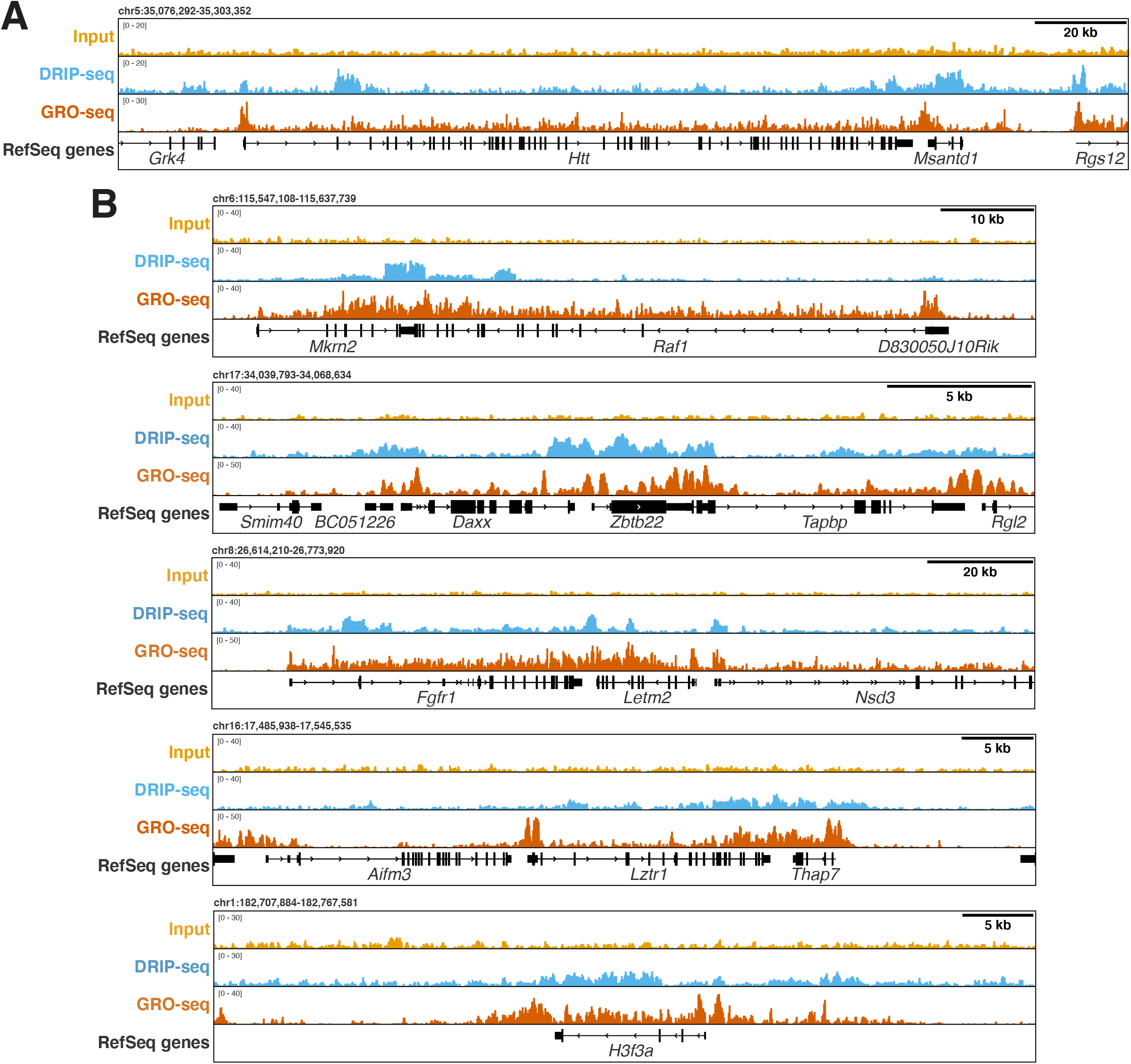
NSPCs show R-loop formation in genes implicated in neurological diseases and brain cancer. **(A)** RPKM-normalized DRIP-seq signal over *Huntingtin* (*Htt*) and **(B)** genes with rearrangements and mutations in human low-grade and high-grade gliomas (*Raf1, Daxx, Fgfr1, Lztr1*, and *H3F3A*). Combined signal from nine DRIP samples and matching input controls from three biological replicates of APH-treated NSPCs is plotted. RPKM-normalized NSPC GRO-seq signal shows transcriptional activity of the genes. RefSeq genes are shown in black.

